# C subunit of the ATP synthase is an amyloidogenic channel-forming peptide: possible implications in mitochondrial pathogenesis

**DOI:** 10.1101/2020.01.16.908335

**Authors:** Giuseppe Federico Amodeo, Brenda Yasie Lee, Natalya Krilyuk, Carina Teresa Filice, Denis Valyuk, Daniel Otzen, Sergey Noskov, Zoya Leonenko, Evgeny V. Pavlov

## Abstract

The c subunit is an inner mitochondrial membrane (IMM) protein and is an integral part of the F_0_ complex of the ATP synthase. Under physiological conditions, this short 75 residue-long peptide folds into an α-helical hairpin and forms oligomers spanning the lipid bilayer. In addition to its physiological role, the c subunit has been proposed as a key participant in stress-induced IMM permeabilization by the mechanism of calcium-induced permeability transition. However, the molecular mechanism of the c subunit participation in IMM permeabilization is not completely understood. Here we used fluorescence spectroscopy, atomic force microscopy and black lipid membrane methods to gain insights into the structural and functional properties of c subunit protein that make it relevant to mitochondrial toxicity. We discovered that c subunit is an amyloidogenic peptide that can spontaneously fold into β-sheets and self-assemble into fibrils and oligomers. C subunit oligomers exhibited ion channel activity in lipid membranes. We propose that the toxic effects of c subunit might be linked to its amyloidogenic properties and are driven by mechanisms similar to those of neurodegenerative polypeptides such as Aβ and α-synuclein.

## Introduction

The c subunit of ATP synthase is a short (75 residue long) peptide whose sequence is highly conserved from bacteria to humans. In eukaryotes, this highly hydrophobic peptide is a transmembrane α-helical “hairpin” localized in the mitochondrial inner membrane where it assembles into oligomers of 8 to 16 units depending on the species. Under normal physiological conditions oligomers of the c subunit are an integral part of the F_0_ complex of the ATP synthase where they form the c-ring that is the main constituent of the rotor^1^.

It has recently been proposed by several groups that, in addition to its normal physiological function, the c subunit may play a critical role in mitochondrial pathology by participating in calcium-induced permeability transition (PT)^2–4^. PT is a phenomenon of increased inner mitochondrial membrane (IMM) permeability by the mechanism of opening of the PT pore in response to toxic levels of reactive oxygen species and/or calcium^5,6^. PT is believed to be a major contributor to cell and tissue damage under conditions of acute stress, including ischemia-reperfusion injury. While multiple experimental studies support the involvement of the c subunit in PT, the molecular basis of its participation remains poorly understood and is a subject of considerable debate^7–10^. Indeed, fractions containing c subunit extracted from mitochondria exhibit channel activity in model lipid bilayers^2,4,11^ suggesting the possibility of its direct participation in PT through the formation of an ion conducting pore. However, in its physiological “c-ring” assembly c subunit is not expected to allow ion flux due to the highly hydrophobic environment within its lumen^12^. We hypothesized that ion conducting capability of c subunit might involve conformational rearrangements and/or assemblies not present in basal conditions. Here we investigated this possibility by testing whether the c subunit protein could form channels in model lipid bilayers. To exclude potential contributions of other biological compounds, we used a synthetic c subunit protein. We combined a structural and functional approach to characterize the peptide and test its channel forming properties. We found that the c subunit is an amyloidogenic peptide that can fold into a β-sheet conformation by forming either fibrils or oligomers in a calcium dependent manner. We observed that the oligomers can form stable pores in model lipid bilayers. We propose that in addition to its native role in the ATP synthase, c subunit can exert its pathological action on mitochondria caused by misfolded oligomers. We hypothesize that the mechanism of c subunit toxicity on mitochondrial membranes might occur through similar structural rearrangements that have been described for other amyloidogenic polypeptides including Aβ (Alzheimer’s Disease) and α-synuclein (Parkinson’s Disease and other synucleinopathies)^13^.

## Materials and methods

### Chemicals

Synthetic peptides with free termini were purchased from LifeTein Inc. (Somerset, NJ, USA) with a purity of 98%.

### Circular Dichroism

CD spectra of synthetic c subunit were recorded on a Jasco J-1500 spectropolarimeter (Jasco Corporation, Japan) with a 1 mm path length cell. Spectra were recorded in the spectral range of 180-260 nm, with a scan rate of 10 nm/min at 0.5 nm intervals. Data were acquired at 25°C, and 10 scans were averaged for each spectrum. Peptide CD spectra were collected in 2% Genapol and phosphate buffer saline pH 7 at a final concentration of 30 μM.

### BLM recordings

The painting method was used to form phospholipid bilayer using 1,2-diphytanoyl-*sn*-glycero-3-phosphocholine (DiPhPC, Avanti Polar Lipids). Bilayer was formed at the 50 – 100 μm diameters apertures of Delrin cuvettes as previously described^14^. Briefly, the aperture was pre-treated with 25 mg/mL of DiPhPC in decane and allowed to dry. Bilayers were formed using the painting method after filling up the cuvettes with the recording solution (150 mM KCl, 20 mM HEPES, pH 7.4) on both sides of the chamber. Ion currents were measured using standard silver-silver chloride electrodes from WPI (World Precision Instruments) that were placed in each side of the cuvette. Measurements of the conductance of single channels were performed by painting the protein to the *cis* side of the chamber (the side connected to the ground electrode). Spontaneous channel insertion was typically obtained under an applied voltage of 20 mV. Conductance measurements were performed using an eONE amplifier (Elements) with a sampling rate of 10 kHz (809.1 μs interval). Traces were filtered by low-pass Bessel filter at 10 Hz for analyses performed with Origin Pro 8 (OriginLab) and Clampfit software (Molecular devices).

### Thioflavin T assay

Fresh stock solutions of c subunit in 2% Genapol/PBS were prepared at room temperature and transferred into a clear-bottomed 96-well plate (CELLSTAR, GBO, Austria) at a final concentration of 5 μM together with 30 μM thioflavin T. Readings were conducted in triplicates either with or without 1 mM Ca^2+^. The plate was loaded into a Flexstation 3 microplate reader (Molecular Devices, San Jose, CA) and incubated at 37 °C without agitation for 20 hours. The fluorescence was measured at 30-sec intervals, with excitation at 440 nm and with emission at 480 nm.

### AFM imaging

The JPK/Bruker Nanowizard II atomic force microscope was used to image each mica slide with c subunit. Aggregates formed without and with 1 mM CaCl_2_ and adsorbed onto the surface of the mica slide. NCH AFM cantilevers were purchased from NanoWorld (Neuchâtel, Switzerland), designed for non-contact and tapping mode imaging to offer high sensitivity and speed while scanning (320 kHz resonance frequency, 42 N/m force constant, thickness 4 μm, no coating). 5×5 μm images were taken in air in Intermittent Contact mode. A minimum of three samples made were used for statistical analysis, with at least six 5×5 μm images obtained for each sample at a resolution of 2048 pixels, and at least 100 measurements were used for statistical analysis of height and diameters calculated.

### SDS-PAGE

Gradient 4-20% Tris-TGX gels were used. Prior to loading, the samples were incubated for 2 hours at room temperature with and without calcium. Next, samples were treated with 5 volumes of 12% (wt/vol) trichloroacetic acid solution and incubated on ice for 5 min. The pellet was collected by centrifugation (11,000 x *g* for 5 min at 4°C) and dissolved in 2% (wt/vol) SDS-containing loading buffer. Prior to being loaded on an SDS-gel, samples were neutralized by the addition of 1 μl aliquots of 0.5 M NaOH until the color of the bromophenol blue-stained loading buffer turned from yellow to blue and were then heated at 98°C for 10 minutes^15^. The separation was performed by applying a constant current of 30 mA. Finally, the gels were stained with silver.

### Bioinformatic analyses

The secondary structure predictions were performed using the Chou & Fasman secondary structure prediction server (CFSSPS) applying a window of three residues^16,17^.

### Data availability

The data that support the findings of this study are available on request.

## Results

### C subunit spontaneously folds into a β-sheet conformation

To investigate the conformational properties of c subunit, we dissolved lyophilized synthetic peptide in a buffer containing a high concentration of a non-ionic surfactant to mimic a membrane-like environment (2 % Genapol and phosphate buffer saline (PBS) at pH 7). After incubation for 15 to 24 hours, we recorded the peptide’s far-UV circular dichroism (CD) spectrum (Figure 1A). The CD spectrum differs markedly from the clear α-helical features of the c subunit found in native conditions, *i.e*. minima around 209 and 222 nm and a maximum around 193 nm. Instead there is a maximum around 198 nm and a minimum around 226 nm. These values are higher than a canonical β-sheet, which typically shows a minimum around 215 nm and a maximum around 195 nm. This suggests additional contributions from other conformations. A CD-spectrum enabled secondary structure analysis performed with BestSel^18^ software suggested that membrane-inserted c subunit oligomers are comprised of ≈44% of antiparallel β-strands and the remaining ≈56% being either turns or other conformations such as β-bridges, bends or unordered structures. Our results are consistent with a secondary structure prediction based on the primary sequence of the c subunit (Fig. 1B and C). The prediction indicates that while the probability of folding into a β-sheet is uniformly distributed along the sequence, there is a gap between A14 and A50 where the score for the a-helix is very low. These results suggest the possibility that similar to some other amyloidogenic polypeptides, if the c subunit is not folded into its native conformation by a functional protein folding machinery it might enter a misfolding pathway that leads to β-sheets.

**Figure 1.**
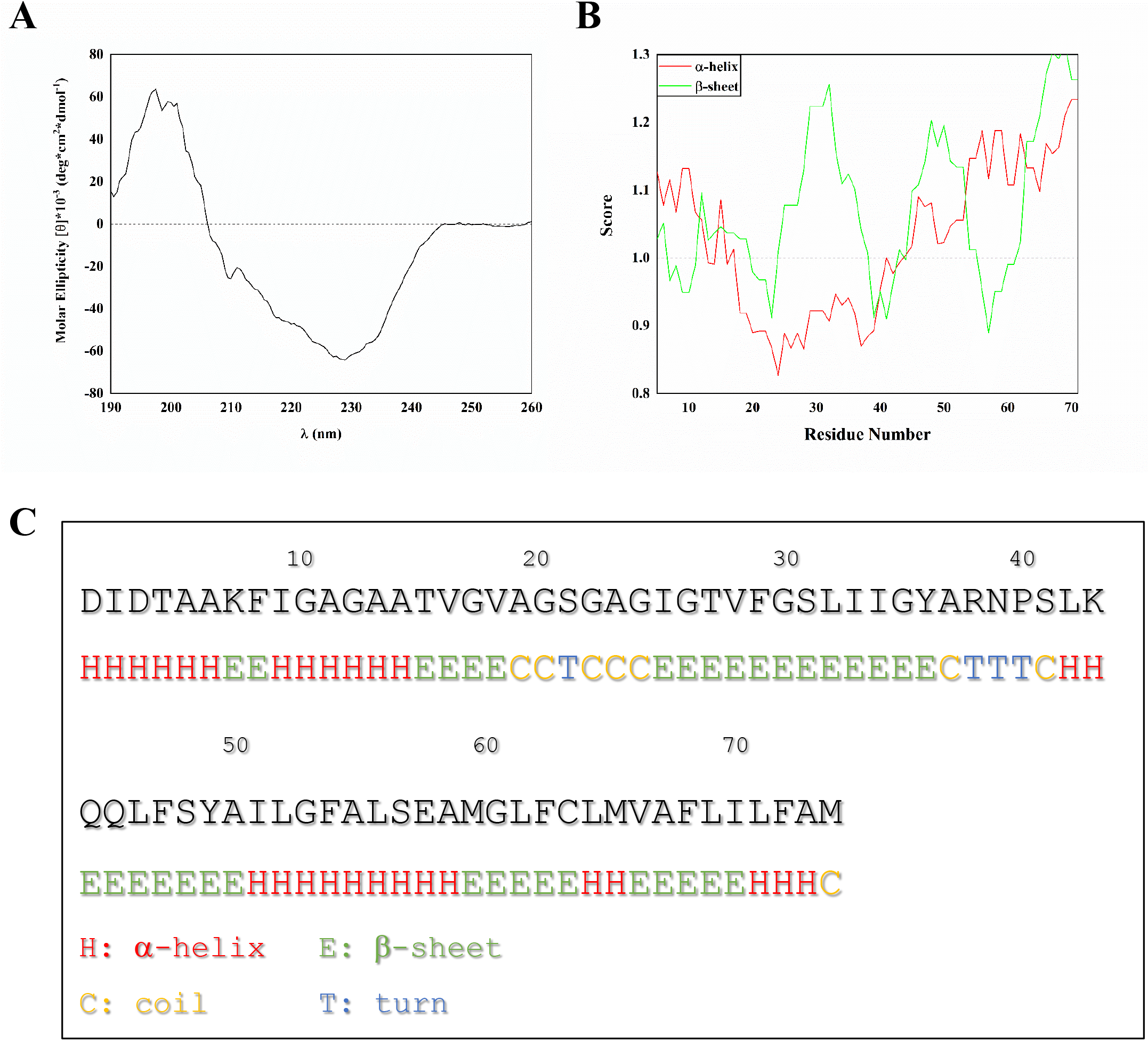
C subunit forms β-sheet structures. (A) CD spectrum of the 50 μM c subunit in 2% Genapol and PBS; (B) Bioinformatic prediction of the probability score of the secondary structure of the c subunit based on its primary sequence according to the Chou & Fasman method; (C) Most probable predicted structure per individual residue.

### Fibrillation and Ca^2+^-induced oligomerization of the c subunit

Next we tested if, similarly to other amyloidogenic peptides, the c subunit in β-sheet conformation could form aggregates. We therefore monitored the formation of cross-β aggregates using the amyloid-binding dye Thioflavin T (ThT). In the absence of Ca^2+^ ions, there was a rapid rise in ThT fluorescence within 3-4 hours at 37°C, which reached a plateau level around 10-15 hrs after the start (Fig. 2A). The aggregation process was strongly suppressed by 1 mM Ca^2+^ (Fig. 2A). The tendency of the c subunit to form complexes was further confirmed by gel electrophoresis (Fig 2B) which shows that the c subunit preparation contains a number to oligomers with MW ranging from 15 up to 250 kDa. Importantly these assemblies were present in both Ca^2+^ and Ca^2+^-free samples, suggesting that calcium ion inhibits fibril formation rather than inducing oligomerization. To visualize the ultrastructure of these c subunit assemblies, we performed atomic force microscopy (AFM) imaging of c subunit samples pre-incubated for 4 hours at 37°C under conditions identical to the ThT assay and deposited on mica. In agreement with CD and ThT data, AFM images of the c subunit samples prepared in the absence of Ca^2+^ showed densely packed fibril structures. The typical diameter estimated from the height measurements of these fibrils was around 20 nm (22.2±0.9nm) (Fig. 2C). In contrast, the presence of 1 mM Ca^2+^ led to the formation of small oligomers with diameters around 60nm (67.6±1.4nm) and few larger aggregates and, importantly, completely inhibited formation of fibrils (Fig. 2D). These assays demonstrate a critical role for Ca^2+^ in the folding and self-assembly of c subunits into oligomers rather than fibrils.

**Figure 2.**
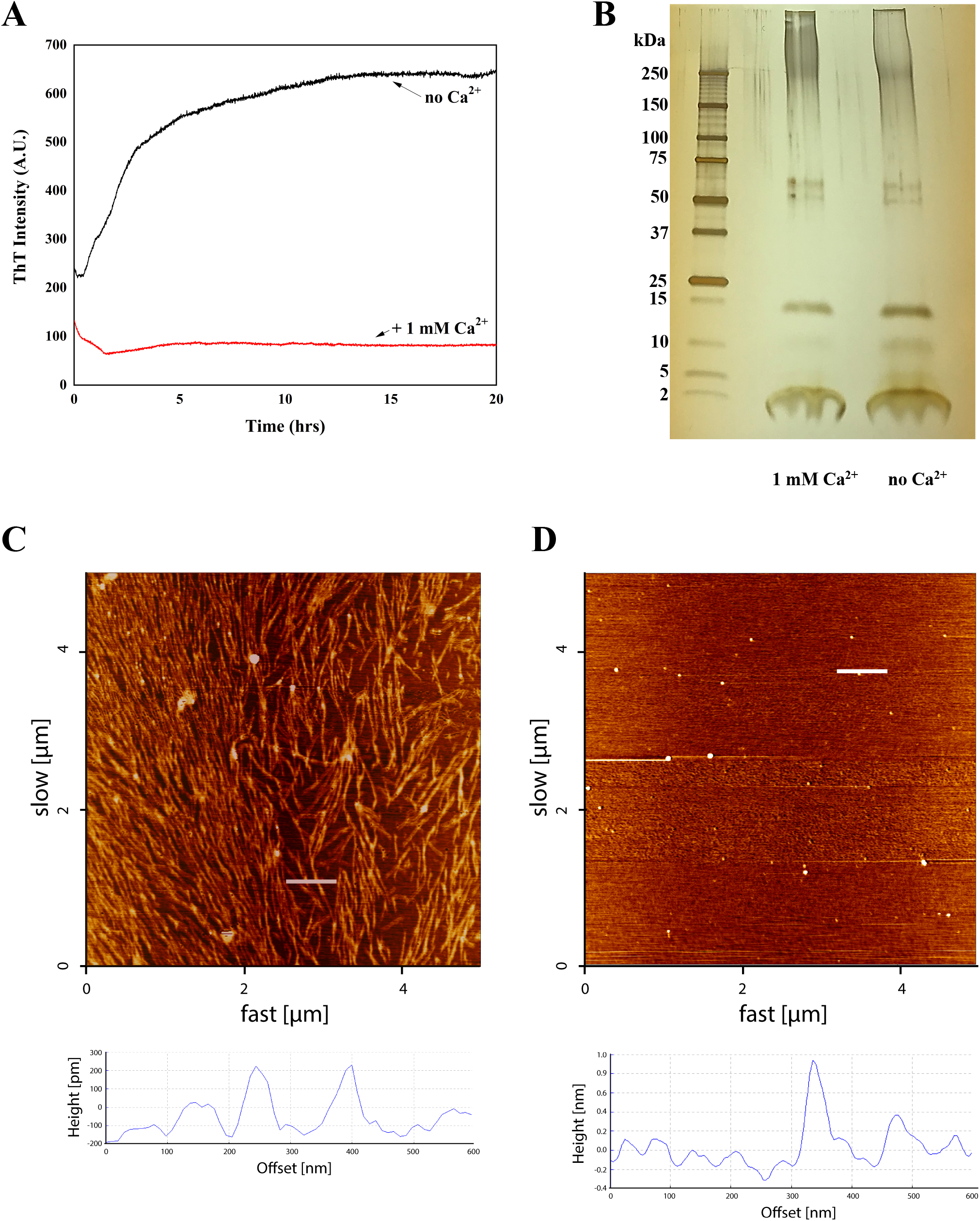
Aggregation properties of the c subunit. (A) Thioflavin T fluorescence spectra showing calcium dependence of c subunit aggregation; (B) SDS PAGE of the c subunit showing presence of oligomeric and monomeric forms; (C) 5×5 μm AFM image of c subunit fibrils after incubation at 37 °C for 4 hours in 2% Genapol and PBS; (D) 5×5 μm AFM image of the c subunit oligomers (arrows) after incubation at 37°C for 4 hours and deposited on mica. Below each image is provided the height profile of the indicated white line.

### Ion channel activity of c subunit oligomers

One of the most critical properties that underlie the toxicity of amyloidogenic peptides is their ability to form ion channels in lipid bilayers^19^. Considering the implications of c subunit in mitochondrial membrane permeabilization and after establishing its amyloidogenic properties we tested its channel forming activity. To do this we reconstituted the preparations described in the previous section into artificial planar lipid bilayers and measured their electrical currents enabled by amyloidogenic peptides by voltage clamping. We observed that both samples prepared with and without Ca^2+^ exhibit robust ion currents. Figure 3A shows representative ion channel behaviour of the c subunit observed in our experiments. These channels show multiple conductance states and were voltage dependent with a tendency to switch to the low conductance states at higher voltages (Fig. 3 B–D). The channel activity was similar in both preparation with slight cation selectivity (P_K_/P_Cl_=6±2, n=8) and an average conductance ranging from 300 to 400 pS. The point distributions of the channel conductances (Fig. 3E) show slightly lower values for the channels from the preparations containing fibrils, but the difference was not statistically significant (p=0.13). The similarity between channel activities of preparations with or without Ca^2+^ suggests that pore forming assembly is not related to the fibrils but rather to oligomers which are present in both samples (Fig 2B). This behaviour is consistent with other known channel-forming amyloidogenic peptides^19,20^.

**Figure 3.**
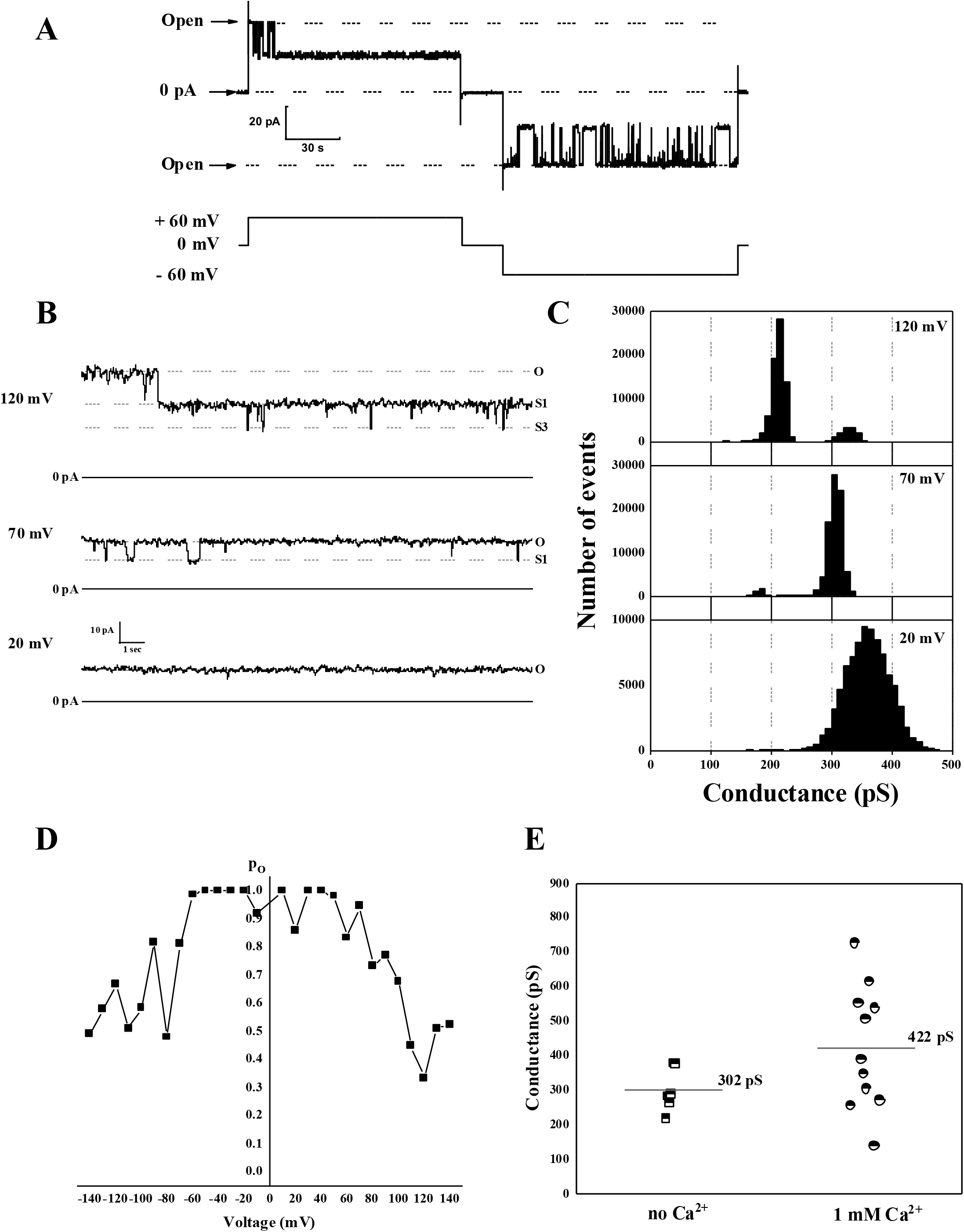
Ion channel activity of c subunit oligomers. (A) Representative current trace of oligomers showing typical channel behaviour with frequent transitions between fully open and lower conductance states; (B) Representative c subunit channel activity at different voltages; (C) all points histogram corresponding to the trace shown at panel B; (D) Voltage dependence of the open probability of the c subunit channel; (E) Channel open state conductance values of the c subunit channels from multiple independent experiments alone (n=5) and in the presence of Ca^2+^ (n=11).

## Discussion

Here we report that synthetic c subunit is an amyloidogenic peptide and its oligomers are capable of forming ion conducting pores in planar lipid bilayers.

A number of previous studies reported that c subunit extracted from mitochondria either as a “c-ring”^2^ or monomer can form channels^4,21^ in lipid bilayers. However, physicochemical properties of c subunit in its native conformation are not consistent with ionophoretic function^12^. It has been argued that the channel forming activity might require the presence of other components that co-purify with the c subunit or post-translational modifications. Our findings indicate that the c subunit alone is sufficient to form ion channels but in order to do so it needs to assemble into oligomers in a β-sheet conformation. We hypothesize that the overall mechanism of the c subunit permeabilization of the inner mitochondrial membrane might be similar to the mechanism of some other amyloidogenic peptides that form β-sheet oligomeric pores^22^. Our work was focused specifically on the synthetic peptide that allowed us to exclude contributions of other biological molecules while focusing on the unmodified c subunit. There is a significant possibility that similar conformational changes might occur *in vivo*. Specifically, it has been established that, similar to other amyloidogenic peptides, c subunit tends to form aggregates both *in vivo* and *in vitro* supporting the notion that it can convert to a toxic form under pathological conditions^21,23^.

The presence of misfolded c subunit peptides could resolve the ongoing controversy regarding PT development through the participation of ATP synthase. None of the currently existing models that require the involvement of ATP synthase in PT provide a mechanistic explanation as to how ion conducting pores can physically form. Our work suggests that such stress induced conformational changes may be the basis of the pore formation. In this scenario the high order rearrangements of the ATP synthase such as F_1_ complex dissociation from F_0_ proposed by Jonas *et al*^2^ or ATP synthase dimerization proposed by Bernardi *et al*^24^ might provide suitable conditions for c subunit transformation to occur.

In summary we report for the first time that c subunit is an amyloid forming peptide and that the ionophoretic properties of c subunit are linked to its amyloidogenic nature and ability to self-assemble into β-sheet oligomers. We propose that toxic forms of misfolded c subunit might play a significant role in cell pathophysiology.

## Acknowledgements

This work was supported by an NIH R01GM115570, United States grant and an American Heart Association, United Sates grant (16GRNT27260229) (to E.V.P.). AFM experiments and instrumentation were supported by NSERC Discovery and CFI grants (to Z.L.) SYN was supported by NSERC Discovery Grant RGPIN-315019, D.E.O. acknowledges support from the Lundbeck Foundation (grant no. R276-2018-671).

## Author contributions

Conception and design of the work: GFA and EP; Acquisition, analysis, and interpretation of data (all authors); Drafted the manuscript GFA and EP; Revised the manuscript (all authors).

